# How quickly can we predict trimethoprim resistance using alchemical free energy methods?

**DOI:** 10.1101/2020.01.13.904664

**Authors:** Philip W Fowler

## Abstract

The emergence of antimicrobial resistance (AMR) threatens modern medicine and necessitates more personalised treatment of bacterial infections. Sequencing the whole genome of the pathogen(s) in a clinical sample offers one way to improve clinical microbiology diagnostic services, and has already been adopted for tuberculosis in some countries. A key weakness of a genetics clinical microbiology is it cannot return a result for rare or novel genetic variants and therefore predictive methods are required. Non-synonymous mutations in the *S. aureus dfrB* gene can be successfully classified as either conferring resistance (or not) by calculating their effect on the binding free energy of the antibiotic, trimethoprim. The underlying approach, alchemical free energy methods, requires large numbers of molecular dynamics simulations to be run.

We show that a large number (N=15) of binding free energies calculated from a series of very short (50 ps) molecular dynamics simulations are able to satisfactorily classify all seven mutations in our clinically-derived testset. A result for a single mutation could therefore be returned in less than an hour, thereby demonstrating that this or similar methods are now sufficiently fast and reproducible for clinical use.

## Introduction

Much of modern medicine relies on being able to prevent and treat bacterial infections. The effectiveness of antibiotics is diminishing since resistance is evolving faster than the rate at which new antibiotics are being developed and brought to market. Antibiotic resistance (AMR) is now accepted as posing a threat to modern medicine requiring urgent and concerted action [1–3]. Clearly activity is required on all fronts, including improving infection control and encouraging the development of new antibiotics. An important part of any solution will be helping clinicians make appropriate treatment decisions by improving the coverage, portability, speed, accuracy and cost of both species identification and antibiotic susceptibility testing (AST). A particularly promising approach is to sequence the genome of any infecting pathogen(s) found in a clinical sample and, by looking up in a catalogue genetic variants known to confer resistance to the action of antibiotics, return a prediction of the effectiveness, or otherwise, of a panel of antibiotics to the clinician [4–8].

Genetic clinical microbiology has been shown to be cheaper, faster and probably more accurate than traditional culture-based clinical microbiology for the AST of tuberculosis [9] and, in addition, facilitates the rapid identification of epidemiological clusters, allowing outbreaks to be rapidly identified. Public Health England adopted whole-genome sequencing for species identification and AST of tuberculosis in 2017 [3, 10] and researchers are validating the approach on other clinically important pathogens. Although catalogues relating genetic variants to resistance phenotype have been carefully and extensively developed, they all share a common weakness: such an approach is fundamentally inferential and so cannot make a prediction when it encounters a genetic variant not present in the catalogue, such as is the case for rare genetic mutations. *Predictive* methods are therefore needed to give the clinician some information about the likely effectiveness of a drug in treating an infection whilst, at the very least, they wait for the clinical sample to be cultured and tested in the traditional manner [11].

Trimethoprim (TMP) is a competitive inhibitor of *S. aureus* dihydrofolate reductase (DHFR, Fig. 1a), an enzyme in the essential folic acid pathway encoded by the chromosomal gene *dfrB*. It is usually administered with sulfamethoxazole, which inhibits another protein in the bacterial folic acid pathway, and the combination is used to treat urinary tract and soft tissue infections. Predicting resistance to trimethoprim is a good test of any novel AMR predictive method since there exists a large amount of structural, biophysical and clinical data and the most common mutation that confers resistance to trimethoprim is F99Y (Fig. 1b), which is a comparatively small mutation and therefore is a stringent test of any predictive method.

**Figure 1:**
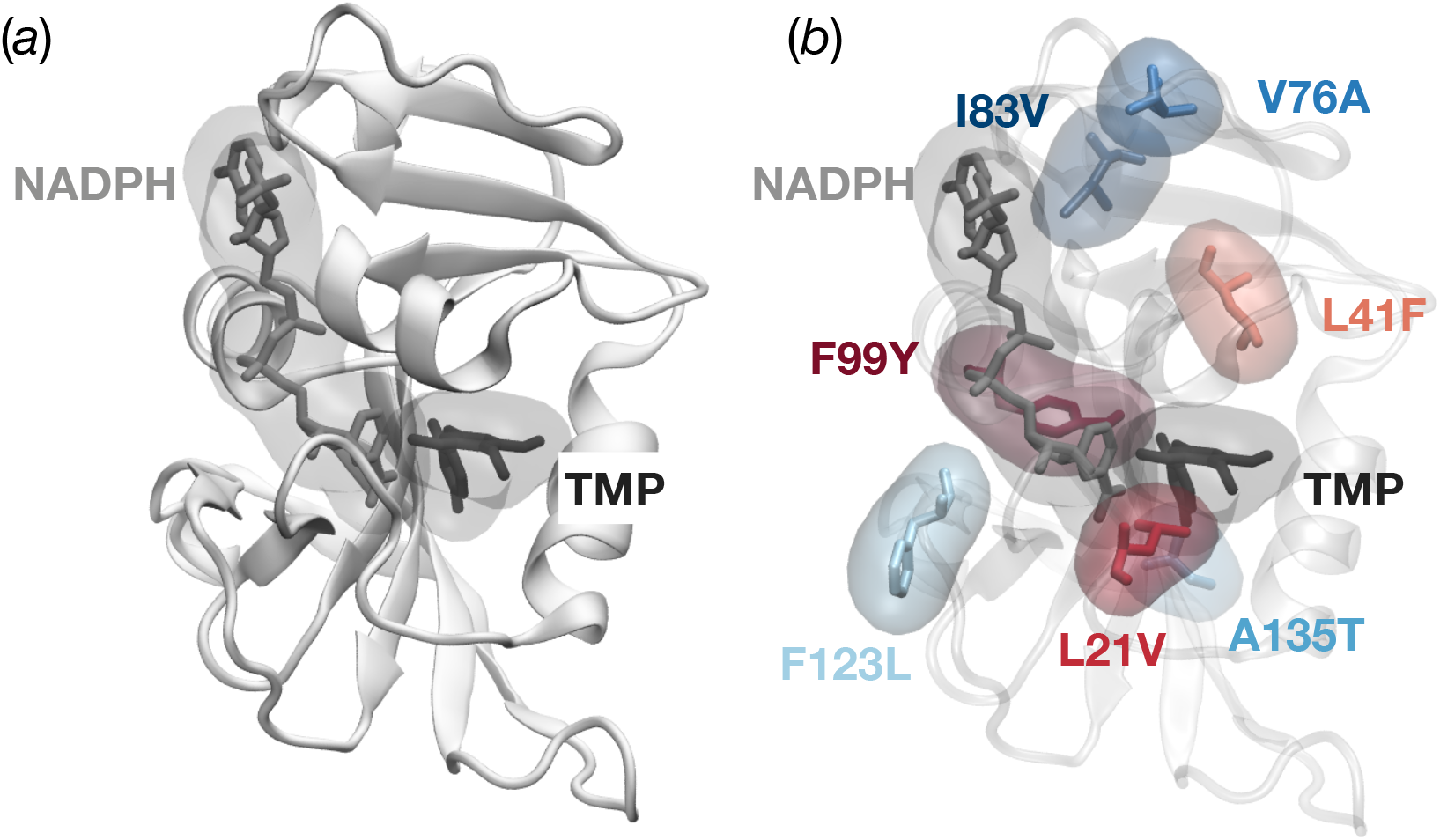
The structure of *S. aureus* DHFR [13] showing (*a*) the overall topology and the trimethoprim (TMP) binding site and (*b*) the location of the seven mutations studied. The three mutations that confer resistance are coloured in different shades of red, whilst the four mutations that have no clinical effect on the action of trimethoprim are coloured in different shades of blue.

Since DHFR is essential, our hypothesis is that non-synonymous protein mutations confer resistance by reducing how well the antibiotic, but not the natural substrate (dihydrofolic acid, DHA), binds. This reduces the problem to calculating how the binding free energy of the drug (ΔΔ*G*_*tmp*_) changes upon introducing the protein mutation. If the mutation reduces the binding free energy below a pre-determined threshold, then one predicts that the mutation confers resistance. Given the subtle nature of the mutations involved and the small size of this protein, we assume that only alchemical free energy methods which are derived from classical statistical mechanics will be sufficiently accurate and precise. It has been previously shown that such methods can be successfully employed to predict the effect of individual amino acid mutations on the action of trimethoprim [12]. By applying simple kinetic theory to clinically observed minimum inhibitory concentrations of trimethoprim for resistant and susceptible samples the previous study was able to establish that for a mutation to confer resistance, ΔΔ*G*_*tmp*_ ≥ 0.8 kcal/mol.

For such a method to be deployed clinically it must be both fast and consume as little computational resource as possible. Whilst broadly successful, the previous study required 32,344 molecular dynamics simulations to be run, yielding a total of 8.1 *μ*s. At the time of writing, one can simulate about 10 ns per day of DHFR using 4 computer cores slaved to a single consumer-grade graphics processing unit (GPU). The calculations underlying a single prediction therefore would require 9,720 CPU hours and 2,430 GPU hours which, although feasible, is still too large for routine use.

Any method must also meet the thresholds for accuracy and reproducibility as laid out by the existing international standards for new antibiotic susceptibility testing methods [14]. The relevant criteria are the very major discrepancy (VMD) and major discrepancy (MD) rates. The former is defined as the number of samples that are classified as susceptible by the method under test which the reference method determined as being resistant as a proportion of the total number of resistant samples and, to pass, VMD ≤ 3%. The definition of the major discrepancy rate is similar but inverted, i.e. the number of samples incorrectly interpreted as resistant that are susceptible. Again, to pass, MD ≤ 3%.

In this paper we shall examine how reducing the computational resource allocated to the calculations affects the qualitative prediction of antibiotic resistance and its reproducibility and thereby answer the question:*“just how quickly can we reliably predict the effect of a mutation in* dfrB *on the action of trimethoprim?”*. The answer to this question will guide whether it is yet feasible to consider deploying this kind of approach clinically.

## Results

### Datasets

A previous study calculated 32 independent values of how the binding free energies of both the antibiotic, TMP, (ΔΔ*G*_*tmp*_) and the natural substrate, DHA, (ΔΔ*G*_*dha*_) changed for each of seven mutations [12]. Here we shall focus solely on the effect of the mutations on the binding free energy of the antibiotic (ΔΔ*G*_*tmp*_). Three of the mutations (F99Y, F99Y/L21V, L41F) are known to confer resistance (Fig. 1b) whilst the remaining four (F123L, A135T, V76A, I83V) have no clinical effect on the action of TMP [15]. Each free energy (ΔΔ*G*_*tmp*_) required 4 alchemical free energies to be calculated (Fig. S1). Two of those energies (Δ*G*_1_ and Δ*G*_6_) were themselves the sum of three further alchemical free energies, hence 8 free energies were required in total. Each free energy in turn necessitated between 8 and 16 molecular dynamics (MD) simulations, each at a different value of the progress parameter, *λ*. By the standards of the field, all of the MD simulations were short, at just 250 ps in duration. We call this existing collection of simulations Set1 (Table 1).So that we may assess whether the calculations are converged, we first extended the simulations underlying ten of the 32 free energies by an order of magnitude, i.e. to 2.5 ns (Set2, Table 1). To allow us to further assess the convergence of the calculations, we ran an additional five calculations for the F99Y mutation using simulations 25 ns long; we call this Set3 (Table 1).

**Table 1:** We took a published set of thirty two values of ΔΔ*G*_*tmp*_ per mutation (Set1) and extended the simulations underlying ten of these values by an order of magnitude creating a second set (Set2). A further five values of ΔΔ*G*_*tmp*_ were calculated for the most commonly observed mutation associated with resistance, F99Y, using simulations two orders of magnitude longer than Set1 (i.e. 25 ns). Note therefore that Set2 is a subset of Set1 since the first 250 ps of all the simulations in Set2 appear also in Set1. Since an additional five values of ΔΔ*G*_*tmp*_ were calculated for F99Y mutation (Set3), Sets 1 & 2 for this mutation contain 5 more members than the other mutations with 37 and 15 members, respectively.

| Name | Number of mutations | Number of values of $\Delta\Delta G_{tmp}$ per mutation | Simulation duration (ns) |
| --- | --- | --- | --- |
| Set1 | all | 32 | 0.25 |
| Set2 | all | 10 | 2.5 |
| Set3 | F99Y only | 5 | 25 |

### Assessing the convergence of the free energy calculations

First let us consider how extending the duration of the molecular dynamics simulations affects the accuracy and precision of how F99Y alters the binding free energy of trimethoprim, ΔΔ*G*_*tmp*_. For simplicity, in this and all subsequent calculations, the first half of each trajectory is discarded to remove transients. The mean and standard deviation of ΔΔ*G*_*tmp*_ is plotted as a function of how much data are included, as described by the simulation duration, *t* (Fig. 2). During the first few tens of picoseconds (Set1) the variance decreases, as one might expect, with the mean remaining approximately constant up to 0.25 ns and in agreement with the published thermodynamic data for this mutant measured using isothermal titration calorimetry (ITC) [13, 16–19]. Swapping to Set2 allows us to examine the behaviour in the range 0.25 ns *< t* ≤ 2.5 ns at the cost of a reduced number of values of ΔΔ*G*_*tmp*_ from which to calculate statistics (n=15 v n=37, Table 1). Here, the mean value of ΔΔ*G*_*tmp*_ rises and then falls slightly, however, the variance is observed to gradually *increase*. Again, it agrees with the published ITC data [13, 16–19]. Finally, moving into the range 2.5 ns *< t* ≤ 25 ns using Set3 we observe the mean value of ΔΔ*G*_*tmp*_ to fluctuate between 0 and 2 kcal/mol whilst the variance remains similar, or increases slightly. The net result is here there is only intermittent agreement with the published ITC data [13, 16–19]. Note that only 5 simulations are contributing to the statistics in this last regime (Table 1) and therefore one is least confident about the observed trends.

**Figure 2:**
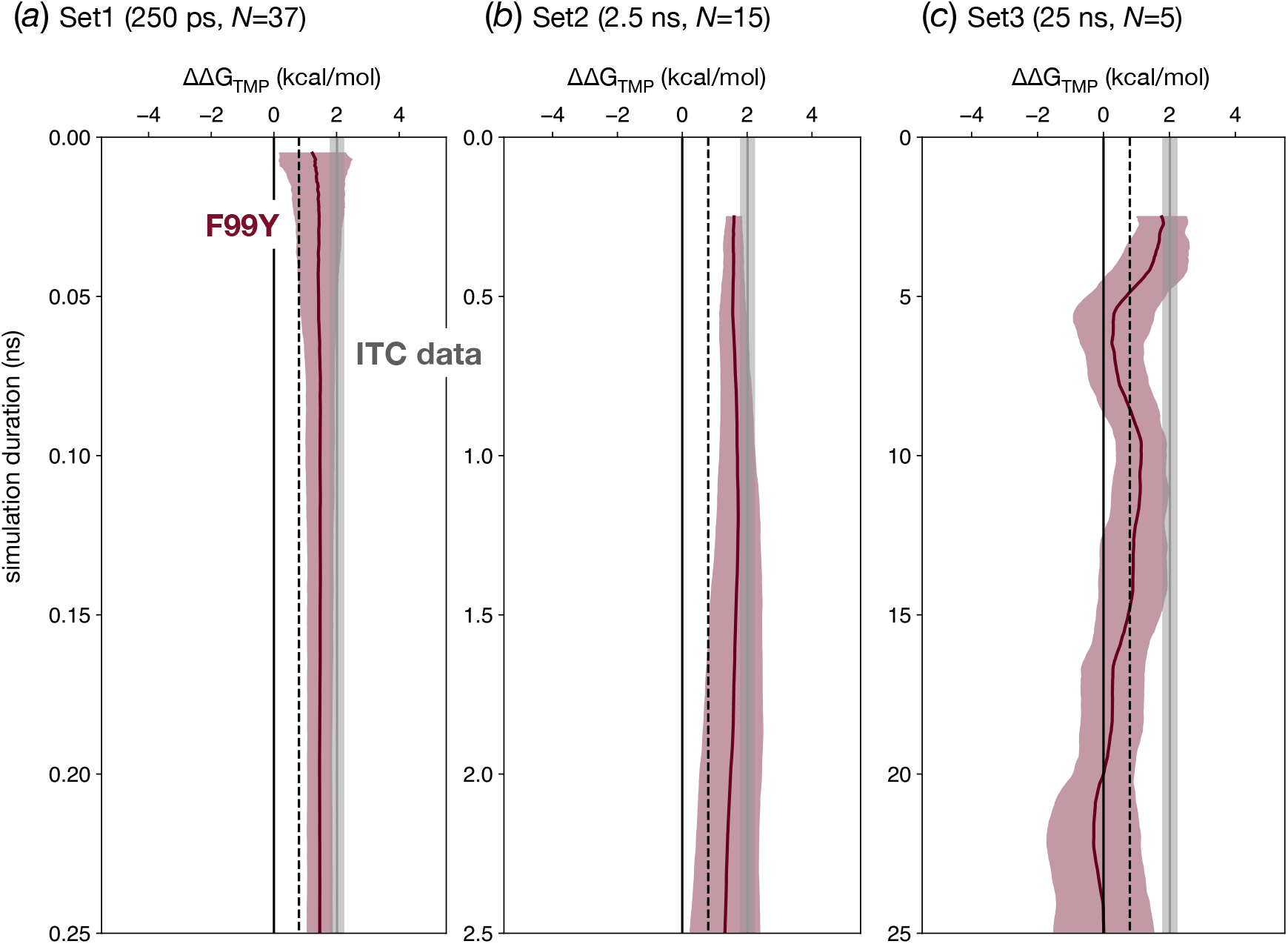
At short simulation durations, the value of ΔΔ*G*_*tmp*_ is in agreement with the results of published isothermal titration calorimetry (ITC) data [13, 16–19], however, in the region 2.5 *< t* ≤ 25*ns* there is scant agreement due to the large deviations in the mean of ΔΔ*G*_*tmp*_. For each of the three sets of simulations, the mean and standard deviation of the values of ΔΔ*G*_*tmp*_ are plotted as a function of the simulation duration, *t*. The dotted line is the threshold (0.8 kcal/mol) above which a mutation would be clinically classified as resistant [12] and the shaded grey area is the 95% confidence limit for the four published values of ΔΔ*G*_*tmp*_ for the F99Y mutation [13, 16–19].

The thermodynamic cycle (Fig. S1) describes how ΔΔ*G*_*tmp*_ can be decomposed into four separate free energies,

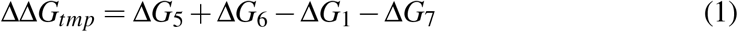

where Δ*G*_1_ and Δ*G*_6_ are the free energies of introducing the mutation into the protein when trimethoprim is either absent or bound, respectively. To prevent the ligand unbinding from the protein, a simple distance restraint was applied during the Δ*G*_6_ transition. The remaining two free energies, Δ*G*_5_ and Δ*G*_7_, estimate the cost of removing this restraint from the mutant and wildtype protein respectively. By considering Set1, Set2 & Set3 in turn, the variation of all four of these free energies in the regime 0.05 *< t* ≤ 25 ns is shown in Fig. S2.

Once *t >* 5 ns There is little change in the mean and the standard deviation is small for the two alchemical free energies, Δ*G*_1_ and Δ*G*_6_, and hence they appear to have converged. Interestingly, although when *t* ≤ 5 ns the means of both values drift (especially in Set2), they appear partially correlated, such that the mean of the difference, as prescribed by Equation 1, varies less than its components. Note also that at small values of *t* (Set1), the variance in Δ*G*_1_ (apo) is larger than Δ*G*_6_ (complexed), which might be expected given the experimental structure had trimethoprim bound. By contrast, the variances of the free energies of removing the restraints (Δ*G*_5_ and Δ*G*_7_) are small at the shorter simulation durations (Set1) and grow in Set2 before in Set3 approaching 4 kcal/mol whilst the mean fluctuates by around 2 kcal/mol.

The observed reduction and then subsequent growth in the variance of ΔΔ*G*_*tmp*_ for the F99Y mutation observed in Fig. 2 is hence due to two different effects; at short times (Set1, as previously published [12]) increasing the duration of the simulations reduces the apparent variance in the alchemical free energies, Δ*G*_1_ and Δ*G*_6_, and there appears to be little contribution from the cost of removing the restraints. But as the simulations are extended, the variances of the restraining free energies, Δ*G*_5_ and Δ*G*_7_, increase until it is their variances that are dominating the apparent error of ΔΔ*G*_*tmp*_.

Similar trends are observed for the other six mutations (Fig. 3); the variance in ΔΔ*G*_*tmp*_ decreases with simulation duration before increasing again. Inspecting the four component free energies (Fig. S3 & S4) shows that, with the exception of F123L, the initial decrease in the variance of ΔΔ*G*_*tmp*_ is mainly driven by convergence of the two alchemical free energies, Δ*G*_1_ and Δ*G*_6_, and, like F99Y, any subsequent increase in the variance is mainly driven by the increasing variance of the free energies of removing the restraints (Δ*G*_5_ and Δ*G*_7_). And, as one would expect, these effects are most pronounced for the mutations where the largest number of atoms are being perturbed (L41F, F123L).

**Figure 3:**
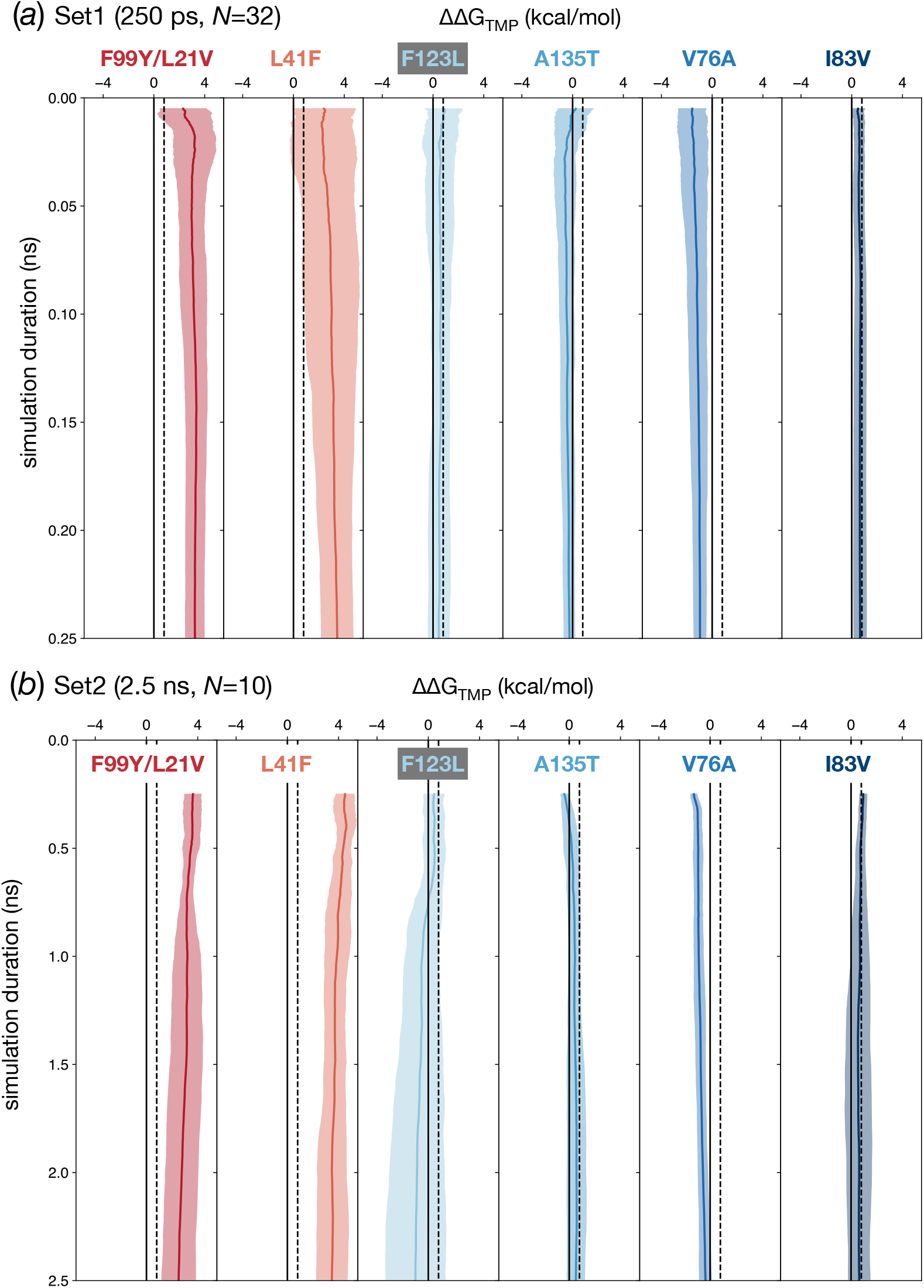
The mean and standard deviations as a function of simulation duration for the other six mutations. As seen in the F99Y mutation (Fig. 2), the variance in ΔΔ*G*_*tmp*_ initially falls as the simulation duration increases, but then rises again. To allow a wide range of simulation durations both (a) Set1 and (b) Set2 were used which contain 32 and 10 values of ΔΔ*G*_*tmp*_ per mutation, respectively. The dotted black line is the resistance threshold [12]: if ΔΔ*G*_*tmp*_ is greater than this value then we predict that it will confer resistance to trimethoprim.

Why is the variance of the cost of removing the restraints increasing with the simulation duration? Since the restraining potential is weaker for high values of the alchemical progress parameter, *λ*, one possible reason might be that after sufficient time the ligand is able to unbind or change conformation. Visually inspecting the simulation trajectories confirms that this is not the case, although it remains possible a more subtle effect is occurring that has not been detected. Alternatively, a more complex set of restraints or a flat-bottomed harmonic potential may behave better, however that is outside the scope of the present work.

In absolute free energy calculations, restraints are required to keep the ligand in the bound state as the ligand is decoupled [20, 21]. For relative binding free energy calculations, as done here, a restraint is only necessary if the ligand departs from the bound conformation during the simulation [20]. To determine retrospectively if restraints are needed for these mutations, five unrestrained 25 ns simulations of each end-state (e.g. F99Y) with trimethoprim bound for each of the seven mutations were run. The ligand did not depart from the binding site in any of the simulations (Fig. S5), strongly suggesting that restraints are not, in fact, required for these calculations. That said, unless one runs a series of unrestrained simulations beforehand to check if the mutation does not cause the ligand to unbind, it remains best practice to include restraints to prevent any unbinding events from occurring.

We conclude that the relative binding free energy calculations shown here are not converged, although we note that some of the component free energies (Δ*G*_1_ and Δ*G*_6_) for some of the mutations show some evidence of converging as *t* → 2.5 ns. Due to the lack of convergence, one must assume that the resulting numerical values of ΔΔ*G*_*tmp*_ are likely inaccurate and that the reported errors are underestimating the uncertainty. We are, however, primarily interested here in how rapidly can we make a *qualitative* prediction of antibiotic resistance and so we shall now turn our attention to this.

### Reducing the simulation duration decreases the sensitivity of the AMR prediction

Let us use Set1 to examine how the method performs when simulations with *very* short durations are used to predict the effect of each mutation on the binding, and thence action, of trimethoprim. We will evaluate the method by comparing its ability to predict the effect of the individual mutations and then, by aggregating the results, calculate the overall sensitivity and specificity and the very major discrepancy (VMD) and major discrepancy (MD) rates.

At a specified simulation duration, *t*, ten values of ΔΔ*G*_*tmp*_ were drawn with replacement from Set1 for the mutation being studied. The mean value of ΔΔ*G*_*tmp*_ and its associated error (95% confidence) were then calculated. If this value was greater than the threshold of 0.8 kcal/mol that defines clinical resistance [12] than the mutation was predicted as conferring Resistance (R); conversely, if this value lay below the threshold, it was predicted as having no effect, i.e. the sample would be Susceptible (S) to trimethoprim. Finally if its confidence limits bracketed the threshold, then it was classified as having an Unknown effect (U). This was repeated 100 times at this value of *t* (Fig. 4), thereby allowing us to examine the reproducibility of the method. By then repeating the process across all mutations for all values of *t*, it also allows us to examine how reducing the simulation duration affects our ability to predict the effect of each mutation on trimethoprim when only using ten values of ΔΔ*G*_*tmp*_ are used.

**Figure 4:**
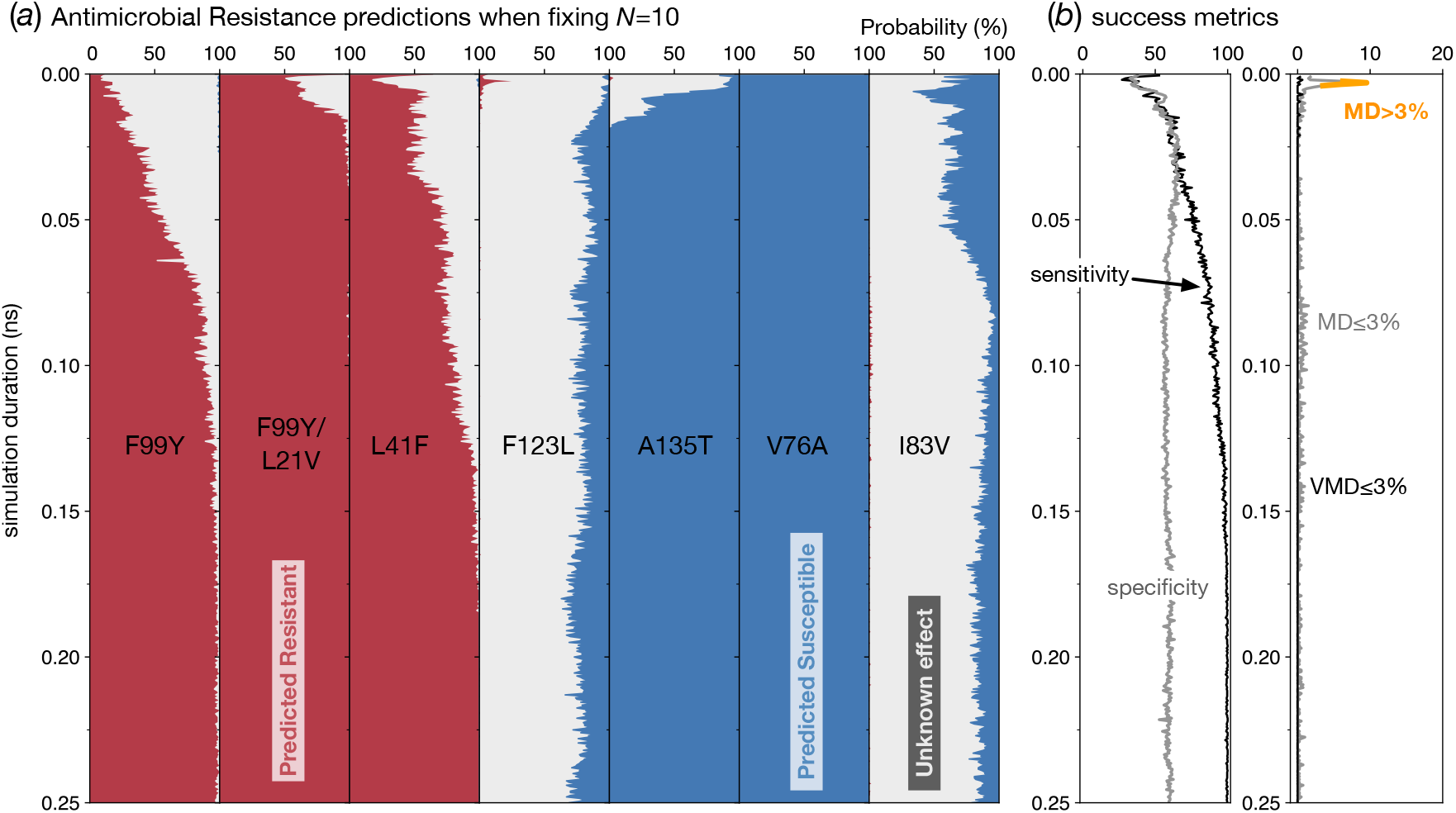
The effect of reducing the simulation duration when using a fixed number of ΔΔ*G*_*tmp*_ values to make a prediction. (*a*) If the number of values of ΔΔ*G*_*tmp*_ used to predict the effect of the mutation on the antibiotic constant (*N* = 10) is kept constant and the simulation duration is decreased, then there is an increase in the proportion of the time where the method cannot make a definitive prediction (i.e. returns ‘Unknown’). Note that the behaviour varies depending on the mutation (e.g. V76A). (*b*) This results in a fall in the overall sensitivity of the method as shorter simulations are used to calculate ΔΔ*G*_*tmp*_. The overall specificity remains approximately constant. (*c*) The very major discrepancy (VMD) rate remains below the required 3% threshold throughout whilst the major discrepancy (MD) rate reaches a maximum of 10% before falling below the threshold after t ~ 0.01ns.

If we allow the method to use all the data (*t* = 250 ps) then it correctly and reproducibly predicts the three mutations clinically associated with trimethoprim resistance (F99Y, F99Y/L21V and L41F). This can be deduced from the earlier graphs of ΔΔ*G*_*tmp*_ (Fig. 2 & 3) since the values for these three mutations and their variances lie above the clinical threshold for resistance. Of the four mutations known to have no effect, the method correctly and reproducibly predicts two to be susceptible (A135T and V76A). The other two are predicted either to be susceptible or to have an unknown effect (F123L and I83V), with the latter more probable than the former.

Now let us consider the effect of reducing the duration of the alchemical simulations. First, and perhaps most importantly, we observe that, across all simulation durations, the method very rarely makes an incorrect definitive prediction i.e. falsely predicts a mutation either to confer resistance when it has no effect (a major discrepancy) or to be susceptible when it confers resistance (a very major discrepancy). The clearest example is when a very short duration is used the F123L mutation is incorrectly predicted to confer resistance with a ~ 25% probability. As a result, we see that as the duration of the simulations is decreased, the probability that the method returns an Unknown prediction rises. The net effect of this is that once the method has returned a definitive prediction (i.e. Resistant or Susceptible) then that is overwhelmingly likely to be correct.

If we combine the results for all the mutations, the resulting sensitivity is high (*t* ≥ 0.5 ns: mean 94.9 %, maximum 100 %, minimum 74.3 %), whilst the specificity varies between 50-60 % (*t* ≥ 0.5 ns: mean 58.6 %, maximum 64.0 %, minimum 53.8 %). Formally since the definitions of sensitivity and specificity assume a binary not a ternary phenotype, the implication of a low specificity is that the method is incorrectly classifying Susceptible samples as Resistant, which is not the case since they are, for the most part, being classified as Unknown. We are therefore perhaps being unduly conservative by including in the denominator cases where an Unknown phenotype has been predicted.

### Using a fixed amount of computational resource

Keeping the number of values of ΔΔ*G*_*tmp*_ contributing to the average constant ensures that the amount of computer resource *increases* as the length of the simulations increase. Perhaps a more helpful question to answer is, if one has a *fixed* amount of computational resource, should one calculate a large number of values of ΔΔ*G*_*tmp*_ using very short simulations, or should one instead calculate a few values of ΔΔ*G*_*tmp*_ using longer simulations? Since using 10 values of ΔΔ*G*_*tmp*_ with *t* = 250 ps (Fig. 4) drawn from Set1 led to an acceptable prediction performance, let us try using 30% of the computational resource those simulations required i.e. the equivalent of 3 values of ΔΔ*G*_*tmp*_ calculated using simulations 250 ps long (which works out at 48 ns of simulation per prediction). Set1 has 32 independent values of ΔΔ*G*_*tmp*_ for each mutation available (37 for F99Y) and hence there is a sufficiently large number of values for bootstrapping in the region 25 *< t <* 250 ps.

Only one of the three resistant mutations (F99Y/L21V) is classified as conferring Resistance throughout (Fig. 5) – the other two have a probability of 20-40% of being predicted having an Unknown effect on trimethoprim. Two of the mutations known to have no effect on trimethoprim (V76A & A135T) are consistently classified as Susceptible, with both having a small probability (5-10%) of being classified Unknown at larger values of *t*. The other two mutations (F123L & I83V) are most likely to be predicted as having an Unknown phenotype, with both consistently having a very small probability of being incorrectly predicted as Resistant and F123L having a high probability of being incorrectly classified as Resistant when *t <* 20 ps. Aggregating these results leads to approximately constant values of the sensitivity (*t* ≥ 0.5 ns: mean 81.2 %, maximum 90.0 %, minimum 71.3 %) and specificity (*t* ≥ 0.5 ns: mean 54.7 %, maximum 69.0 %, minimum 45.2 %), although again we emphasise that by including the cases predicted Unknown in the denominator of the sensitivity and specificity calculations, we are probably being over-conservative and excluding these cases would result in much higher values for the sensitivity and specificity.

**Figure 5:**
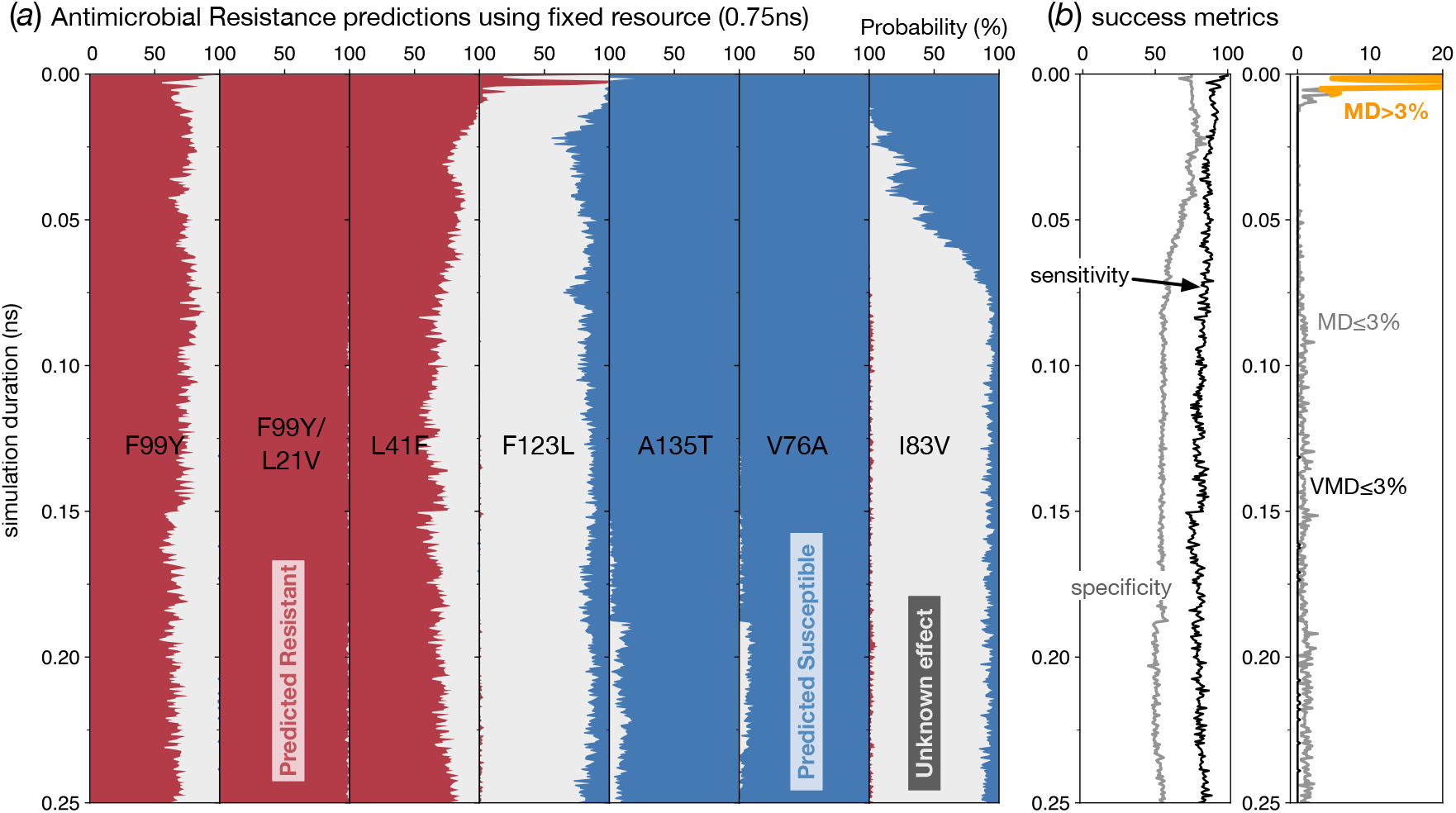
The effect of using a fixed amount of computational resource on prediction. (*a*) If the amount of computational resource used to predict the effect of the mutation on the antibiotic constant (equivalent to N=3 using 250 ps simulations) is kept constant then the number of calculations that can be run increases as the simulation duration is decreased. The method tends to either make a correct definitive prediction (Resistant or Susceptible) or return an Unknown result. (*b*) The greater probability of making an Unknown prediction results in a lower specificity, however the sensitivity remains high. (*c*) The very major discrepancy (VMD) rate remains below the required 3% threshold throughout whilst the major discrepancy (MD) rate reaches a maximum of 25% before falling below the threshold after t ~ 0.01ns.

### How fast could we predict whether a mutation confers resistance to trimethoprim?

The detailed analysis in Fig. 4 & 5 enables us to choose values of (*N, t*) to minimise the time required to make a satisfactory set of predictions, however one does not *a priori* usually have this information and, in any case, it is possible that as yet unseen mutations in *dfrB* could have effects not congruent with our testset of seven mutations. For example, our testset does not contain any mutation that is marginally resistant (which explains why the sensitivity is always greater than the specificity). This analysis therefore should be taken as indicative, however, it is clearly of interest to consider just how rapidly one could predict whether a mutation confers resistance to trimethoprim or not.

We shall choose *N* = 15 and *t* = 50 ps (Fig. 5) since the trajectories are long enough to avoid the observed transients in classification behaviour which affect the major discrepancy rate but are still short enough to run quickly using a consumer grade GPU. Using these parameters the sensitivity and specificity are predicted to be 86.3% and 70%, respectively, with no very major discrepancies (VMD) or major discrepancies (MD). The lack of any VMDs or MDs illustrates again that the method is either returning a correct prediction of Resistant or Susceptible, or is returning a prediction of Unknown. We estimate that, at the time of writing, a single 50 ps trajectory will take ~ 7.5 minutes to complete, assuming the GPU is large enough (or there is more than one on the motherboard) so that all the replicas can run on the same node to permit Hamiltonian replica exchange which requires rapid communication between the replicas. Since we have restricted ourselves in this paper to only calculating ΔΔ*G*_*tmp*_, each value therefore only requires 8 alchemical free energies to be calculated, each in turn requiring (at least) 8 coupled molecular dynamics simulations, making a total of 960 simulations, which is still daunting. If we arbitrarily decide that a prediction must be complete within one hour, then twenty nodes, each with one or more GPUs and 8-16 CPU cores, could run the simulations in 45 minutes. Such a facility could easily be provided by a large research University or a commercial cloud platform. That would leave 15 minutes for setup and analysis: preparing all simulation input files on a single machine would likely create a bottleneck, and hence one would also have to distribute all the setup tasks onto the high performance computer. We conclude that, whilst there are obvious challenges, it is now feasible to predict whether individual mutations confer resistance to an antibiotic using free energy methods and that this can be done fast enough to be clinically useful.

### Prevalence of the studied mutations in the European Nucleotide Archive

The seven different non-synonymous *dfrB* mutations studied here were selected from a relatively small dataset of 501 unrelated *S. aureus* isolates collected from patients in the UK [12, 15]. Our analyses and conclusions depend on our testset of seven mutations being representative of the likely mutations in *dfrB* one might encounter clinically. To estimate how prevalent these mutations are globally, we searched an index of the European Nucleotide Archive [22]. Due to how the index is created, only results for amino acids 11-148 (incl.) were returned, and samples containing multiple amino acid mutations fewer than ten positions apart are unlikely to have been detected. That said, this is as comprehensive a scan of all deposited *S. aureus* genomes as is currently possible and approximately 19,200 *S. aureus* genomes were searched (Table S1). Although F99Y was found in 13.8 % (69) of the clinical isolates, the prevalence in the ENA was only 0.7 % (137), confirming that the clinical dataset was substantially enriched for trimethoprim resistance. Both L41F and L21V F99Y were only detected once in the original clinical dataset (0.2 %). Whilst the double L21V F99Y was not present in the much larger ENA dataset, 421 genomes (2.3 %) containing the L21V mutation without F99Y were detected, however the effect of L21V on its own on the effectiveness of trimethoprim is unknown. The L41F mutation was found in the ENA, but at a very low prevalence (0.02 %, n=3). Of the four mutations, A135T (n=162, 32.3 %), V76A (89, 17.8 %), I83V (8, 1.6 %) and F123L (5, 1.0 %), identified as susceptible in the clinical study [15], only two were found in the ENA: A135T (n=6,578, 34.3 %) and V76A (1,412, 7.4 %). Further examination of the results returned for *k*-mers used to probe codon variation at Val76 and Phe123 showed that comparatively few sets of short-reads in the ENA were identified around these positions, suggesting that some samples were missed since they contained multiple amino acid mutations within the width of the *k*-mer (21 amino acids) or that similar *k*-mers are found in other species and therefore our ability to detect variation at these sites using this method is probably limited.

## Discussion

We conclude that it is now possible to rapidly and reproducibly predict whether individual non-synonymous mutations confer resistance (or not) to trimethoprim, an antibiotic, and we have shown that, for this system at least, it is theoretically possible and practical to make a prediction in less than one hour. This relies on our observation that calculating a large number of values of ΔΔ*G*_*tmp*_ using very short alchemical molecular dynamics simulations allows the seven mutations in our test set to be adequately classified with acceptable sensitivities and specificities and low very major and major discrepancy rates, thereby using an order of magnitude less computational resource than a previous study [12].

The relatively small amount of computation resource required also opens up the possibility of running calculations ahead of time. For example, given a large enough computational resource one could predict the effect of all possible single-nucleotide non-synonymous mutations in a small protein like DHFR, thereby providing a lookup table so that a prediction can be returned instantaneously to the clinician. One can further imagine that this data could be applied to help de-risk lead compounds in drug development for resistance, since it would reveal how a protein could evade the action of a small-molecule inhibitor by mutation.

We note, however, that the values of ΔΔ*G*_*tmp*_ are not yet converged and this appears to be due to the restraints applied to prevent the ligand leaving the binding pocket. Although this invites further investigation, since we have demonstrated that restraints are not required to keep the ligand bound to the protein, it suggests that in future it may be simpler and preferable to run such calculations without restraints, thereby reducing the computational overhead yet further. What is curious is that whilst the values of ΔΔ*G*_*tmp*_ are not yet converged and their precision is likely underestimated, they appear sufficiently accurate to satisfactorily predict whether a mutation confers resistance to trimethoprim or not across a wide range of simulation durations. If the mutation under study has a value of ΔΔ*G*_*tmp*_ that is close to the resistance threshold, however, then quantitative accuracy will be important.

Our conclusions are reliant on the seven mutations that form the testset and we have investigated whether these are representative of the genetic variation one might expect to observe clinically. Since several of mutations were detected at either very low prevalences or not at all in the European Nucleotide Archive, we conclude that our testset is not truly representative and therefore our conclusions may not transfer into the clinic. Clearly the method needs to be tested on additional *dfrB* mutations as well as other antibiotic/protein target combinations in a wide range of pathogenic bacterial before we can make more definitive statements about its applicability.

Although we were able to show that calculating 15 values of ΔΔ*G*_*tmp*_ from alchemical simulations only 50 ps long led to acceptable classification behaviour, this is almost certainly a form of overfitting since we had a free choice of a wide range of combinations and may have simply chosen one that works well for our testset. It will only be possible to gain confidence through applying the method ‘blind’ to other mutations. We have also restricted ourselves here to only considering the effect of the mutation on the binding of the antibiotic; previous work has shown that taking the effect on the natural substrate, DHA, into account changes and may improve the prediction [12]. Also, all of the mutations studied (with the possible exception of the double mutant, F99Y/L21V) are tractable by alchemical methods: it remains to be seen how successful this approach will be for mutations involving a change in electrical charge, that involve a proline, or simply require a large number of atoms to be perturbed.

Our method assumes that all the eight alchemical free energies required to calculate a single value of ΔΔ*G*_*tmp*_ all converge and behave similarly, which we have shown not to be true (Fig. S2-S4). Further work is needed to assess if different types of alchemical transition (e.g. removing the electrical charges from the alchemical atoms) require more or less simulation time and/or numbers of molecular dynamics simulations. It is possible up to another order of magnitude of savings is available through careful dynamic control (i.e. ‘steering’ [23]) of the makeup and number of alchemical free energies run. This will necessarily complicate the calculation of errors which was done here at the level of ΔΔ*G*_*tmp*_ and in future will likely have to be done at the level of each alchemical free energy with errors then added in quadrature.

The translation of genetics into clinical microbiology shows no sign of abating, with the most progress being made using whole-genome sequencing for antibiotic susceptibility testing (AST) of tuberculosis where catalogues of observed genetic variants and their associated effects on different drugs are most advanced [24]. Since this approach is purely inferential, there remains a need to develop predictive methods [11]. Even a low-quality prediction for a single antibiotic may prove useful clinically, since it will be viewed alongside the results for other drugs which often allows nonsensical predictions to be discounted. Such a Bayesian approach has already been shown to improve AST of tuberculosis [11]. In any case, if the prediction ends up affecting the clinical decision, it is likely that the sample would still be sent for culturing and testing.

Although we have focussed primarily on calculating the effect of the mutation on the binding free energy of the antibiotic, there are a range of other methods that could be brought to bear. Machine learning methods, using genetic, structural and/or chemical features, are likely to be sufficiently accurate and also fast [11, 25, 26]. Such methods could be used to screen out mutations that have no effect on an antibiotic, leaving only the marginal cases for computationally more intensive approaches such as proposed here.

One key advantage of our approach we have not discussed in detail is that it rarely makes an incorrect classification. This potentially enables a guided process whereby simulations are run until a definite (Resistant/Susceptible) prediction is returned, saving further computational resource and time. More work needs to be done if this, or other, methods are to be formally certified for use in AST. For example, to pass the relevant international standard [14], not only must the very major and major discrepancy rates be ≤ 3 %, but also there must be a high level of categorial agreement with a reference method. This will necessitate carefully designed and thorough studies in collaboration with clinical microbiology laboratories using standard methods.

Clinical microbiology is built upon a binary paradigm of a sample being Resistant or Susceptible. Converting a quantitative measurement which has a confidence limit into a binary result necessitates a third category (here called Unknown) for those cases that cannot be definitely classified as R or S. This is a subtle point (and one can probably trace its origins back to the Law of the Excluded Middle) and it is nonsensical to deny predictive and experimental methods the option of returning an Unknown result. The Clinical Laboratory and Standards Institute, which provides clinical microbiology standards mainly to the U.S.A, has adopted a ternary system, however, the European Committee on Antimicrobial Susceptibility Testing have only recently begun to introduce such a category into some antibiotic/pathogen combinations [27]. Many of the clinical tools implicitly assume the binary Resistant/Susceptible paradigm and new tools and language need developing, as we have seen here when calculating the sensitivity and specificity of our method.

As computational resource becomes faster and more widespread, the broader field of molecular simulation is gradually moving away from running single simulations towards running large number of replicas [28, 29]. This potentially exposes underlying drawbacks with the molecular dynamics codes and how we, as computational scientists, typically work. For example, when setting up and running thousands of molecular dynamics simulations the time taken to setup a simulation becomes appreciable. Likewise, one must use some kind of object store or file hierarchy to archive all the simulation files. Without progress in these areas, it is possible that the time taken to setup, copy files, queue simulations and retrieve and analyse files could become the limiting factor in speeding up and therefore applying these methods clinically.

In the field of alchemical free energy calculations much attention has understandably been focussed on demonstrating that free energies can be calculated that agree with experimental data to a high degree of precision. A high degree of precision and accuracy is spurious in this application and one might speculate that applying alchemical methods in this way is a sign that the field is maturing. Finally, whilst thermodynamic integration is usually described as an equilibrium method, it is not obvious if this remains true when the duration of an alchemical simulation is only 50 ps. Despite this, it is both illuminating and encouraging to look back over the last thirty plus years on the progress made by the field of alchemical free energy calculations [30] and infer what might be possible in just a few more years.

## Methods

The GROMACS molecular dynamics [31] simulations underlying the alchemical free energy calculations were setup and run as described previously [12] and followed best practice [32], including using pmx [33] to mutate the wild-type structure of DHFR with trimethoprim bound [13]. In brief, the structure of *S. aureus* DHFR with trimethoprim and NADPH bound (PDB:3FRE) [13] was mutated as required by pmx [33] and water and counter-ions added. To provide a range of starting structures to seed the different alchemical free energy calculations, each mutated protein, in both apo and TMP-bound forms, was subjected to a short 2.5 ns equilibration simulation using an integration timestep of 1 fs. At a range of timepoints along each simulation, the mutant sidechain was ‘phased-in’ over 1000 steps by smoothly increasing *λ*, as per the Alchembed procedure [34]. Six different initial structures were created in this way for each mutant, thereby reducing the likelihood that short simulations are correlated. The generalised AMBER forcefield and AMBER ff9SB-ildn [35] were used throughout. Electrostatic forces were calculated by the particle mesh Ewald method using a real space cutoff of 1.2 nm whilst van der Waals interactions were cutoff at 1.2 nm, with a switching function applied from 0.9 nm. The temperature was maintained at 310 K using a Langevin thermostat with a time constant of 2 ps and the pressure held at 1 bar using an isotropic Parinello-Rahman barostat with a compressibility of 4.46 × 10^−5^ bar^−1^ and a time constant of 1 ps.

The alchemical simulations underlying ten of the thirty-two values of ΔΔ*G*_*tmp*_ for each of the seven mutations were extended by an order of magnitude (from 0.25 ns to 2.5 ns). In addition five further values of ΔΔ*G*_*tmp*_ for the F99Y mutation were calculated using simulations two orders of magnitude longer (25 ns). To cope with the resulting very large numbers of MD simulations, all data was stored in a file hierarchy and tagged using the datreant Python module [36]. All simulation data was then parsed and alchemical free energies were calculated as a function of simulation time *t* using a purpose-written Python class. In all cases, the first half of the available data were discarded and hence the free energy at time *t* was calculated by thermodynamic integration using the energies in the interval (*t/*2, *t*]. Using the simpler method of thermodynamic integration was possible because, as reported previously [12], there were no significant differences in values of Δ*G* calculated by either this method or the more complex multi-state Bennett acceptance ratio estimator. This yielded large tables of alchemical free energies for Set1, Set2 and Set3 and each was stored as a Pandas dataframe [37]. The resulting values of ΔΔ*G*_*tmp*_ were then calculated and stored. These dataframes can be found in the Supplemental Information. Bespoke Python code then read these tables and applied the bootstrapping process described in the main body of the paper to produce the Figures. Standard errors were calculated in the usual way and converted to a 95% confidence interval using the appropriate t-statistic for the number of values. All graphs were plotted using Matplotlib [38] and all protein images were rendered using VMD [39]. The BIGSI search index for microbial genomes [22] is interrogated using a *k*-mer where *k* ≥ 61 bases. Since searching the index for amino acid mutations involves permuting the bases in a triplet, leading to 80 different variations, we wrote a Python module, called `pygsi`, that automatically interrogates the index using a 63-mer constructed with 20 base pairs flanking the codon of interest [40]. Once all the permutations have been tried, the code moves on to the next codon.

## Supporting information

Supplemental Set1

Supplemental Set2

Supplemental Set3

## Author contributions

PWF designed the study, ran the simulations, analysed the data and wrote the manuscript.

## Acknowledgements

The author is grateful to the organisers of the BioExcel Alchemical Free Energy workshop held in Göttingen in May 2019 during which the idea behind this paper was developed.

## Data Accessibility

The three tables containing all the values of ΔΔ*G*_*tmp*_ for the different mutations used to construct the figures in this paper are provided as Supplemental Files in the CSV format and are described in the Supplemental Information. The results of searching the BIGSI bacterial genomic index are provided in Table S1.

## References

1. O’Neill J (2016) Tackling Drug-Resistant Infections Globally: Final Report and Recommendations. Technical report.

2. Davies SC (2013) Annual Report of the Chief Medical Officer - Vol 2. Technical report, Department of Health, UK Government.

3. Public Health England (2018) Tuberculosis in England.

4. Didelot X, Bowden R, Wilson DJ, Peto TEA, Crook DW (2012) Nat Rev Genetics 13:601–12.

5. Köser CU, Ellington MJ, Cartwright EJP, Gillespie SH, Brown NM, Farrington M, Holden MTG, Dougan G, Bentley SD, Parkhill J, Peacock SJ (2012) PLoS pathogens 8:e1002824.

6. Ellington M, Ekelund O, Aarestrup F, Canton R, Doumith M, Giske C, Grundman H, Hasman H, Holden M, Hopkins K, Iredell J, Kahlmeter G, Köser C, MacGowan A, Mevius D, Mulvey M, Naas T, Peto T, Rolain JM, Samuelsen Ø, Woodford N (2017) Clinical Microbiology and Infection 23:2–22.

7. Tagini F, Greub G (2017) Eur J Clin Micro Infect Dis 36:2007–2020.

8. Balloux F, Brønstad Brynildsrud O, van Dorp L, Shaw LP, Chen H, Harris KA, Wang H, Eldholm V (2018) Trends in Microbiology 26:1035–1048.

9. Pankhurst LJ, del Ojo Elias C, Votintseva AA, Walker TM, Cole K, Davies J, Fermont JM, Gascoyne-Binzi DM, Kohl TA, Kong C, Lemaitre N, Niemann S, Paul J, Rogers TR, Roycroft E, Smith EG, Supply P, Tang P, Wilcox MH, Wordsworth S, Wyllie D, Xu L, Crook DW (2016) Lancet Resp Med 4:49–58.

10. Walker TM, Cruz ALG, Peto TE, Smith EG, Esmail H, Crook DW (2017) Lancet Infec Disease 17:359–361.

11. Brankin AE, Fowler PW (2019) ACS Central Science 5:1312–1314.

12. Fowler PW, Cole K, Gordon NC, Kearns AM, Llewelyn MJ, Peto TE, Crook DW, Walker AS (2018) Cell Chemical Biology 25:339–349.e4.

13. Oefner C, Bandera M, Haldimann A, Laue H, Schulz H, Mukhija S, Parisi S, Weiss L, Lociuro S, Dale GE (2009) J Antimicrobial Chem 63:687–698.

14. International Organization for Standardization (2007) ISO 20776-2: Clinical laboratory testing and in vitro diagnostic test systems. Technical report, International Standards Organization.

15. Gordon NC, Price JR, Cole K, Everitt R, Morgan M, Finney J, Kearns AM, Pichon B, Young B, Wilson DJ, Llewelyn MJ, Paul J, Peto TEA, Crook DW, Walker AS, Golubchik T (2014) J Clin Microbiol 52:1182–91.

16. Pires DEV, Blundell TL, Ascher DB (2015) Nuc Acid Res 43:D387–D391.

17. Dale GE, Broger C, D’ Arcy A, Hartman PG, DeHoogt R, Jolidon S, Kompis I, Labhardt AM, Langen H, Locher H, Page MG, Stüber D, Then RL, Wipf B, Oefner C (1997) J Mol Biol 266:23–30.

18. Frey KM, Georgiev I, Donald BR, Anderson AC (2010) Proceedings of the National Academy of Sciences 107:13707–13712.

19. Frey KM, Viswanathan K, Wright DL, Anderson AC (2012) Antimicrob Agent Chemo 56:3556–3562.

20. Mey ASJS, Allen B, Macdonald HEB, Chodera JD, Kuhn M, Michel J, Mobley DL, Naden LN, Prasad S, Rizzi A, Scheen J, Shirts MR, Tresadern G, Xu H (2020) Best Practices for Alchemical Free Energy Calculations. arXiv:2008.03067

21. Mobley DL, Chodera JD, Dill KA (2006) J Chem Phys 125:084902.

22. Bradley P, den Bakker HC, Rocha EPC, McVean G, Iqbal Z (2019) Nature Biotechnology 37:152–159.

23. Fowler PW, Geroult S, Jha S, Waksman G, Coveney PV (2007) J Chem Theory Comput 3:1193–1202.

24. The CRyPTIC Consortium, 100000 Genomes Project (2018) New Eng J Med 379:1403–1415.

25. Carter JJ, Walker TM, Walker AS, Whitfield MG, Morlock GP, Peto TE, Posey JE, Crook DW, Fowler PW (2019) bioRxiv doi:101101/518142.

26. Aldeghi M, Gapsys V, de Groot BL (2019) ACS Central Science 5:1468–1474.

27. Davies TJ, Stoesser N, Sheppard AE, Abuoun M, Fowler P, Quan TP, Griffiths D, Vaughan A, Morgan M, Phan HTT, Jeffery KJ, Andersson M, Ellington MJ, Ekelund O, Mathers AJ, Robert A, Woodford N, Crook DW, Peto TEA, Anjum MF, Walker AS (2020) Antimicrob Agent Chemo 64:1–16.

28. Knapp B, Ospina L, Deane CM (2018) Journal of Chemical Theory and Computation 14:6127–6138.

29. Bhati AP, Wan S, Hu Y, Sherborne B, Coveney PV (2018) Journal of Chemical Theory and Computation 14:2867–2880.

30. Bash P, Singh U, Brown F, Langridge R, Kollman P (1987) Science 235:574.

31. Abraham MJ, Murtola T, Schulz R, Páll S, Smith JC, Hess B, Lindahl E (2015) SoftwareX 1-2:19–25.

32. Klimovich PV, Shirts MR, Mobley DL (2015) J Comp Aided Mol Des 29:397–411.

33. Gapsys V, Michielssens S, Seeliger D, de Groot BL (2015) J Comp Chem 36:348–54.

34. Jefferys E, Sands ZA, Shi J, Sansom MSP, Fowler PW (2015) J Chem Theo Comp 11:2743–2754.

35. Lindorff-Larsen K, Piana S, Palmo K, Maragakis P, Klepeis JL, Dror RO, Shaw DE (2010) Proteins 78:1950–8.

36. Dotson DL, Seyler SL, Linke M, Gowers RJ, Beckstein O (2016) In Proc 15th Python Sci Conf, edited by S Benthall, S Rostrup, 51–56.

37. McKinney W (2010) In Proceedings of the 9th Python in Science Conference, edited by SvdW Millman, Jarrod, 51–56.

38. Hunter JD (2007) Computing in Science & Engineering 9:90–95.

39. Humphrey W, Dalke A, Schulten K (1996) J Mol Graph 14:33–38.

40. Fowler PW (2017). pygsi: a Python class to interrogate BIGISI. doi: 10.5281/zenodo.1407085. https://github.com/philipwfowler/pygsi.

