## Supplemental Material for "How quickly can we predict trimethoprim resistance using alchemical free energy methods?"

Philip W Fowler[†1,2]

[1]Nuffield Department of Medicine, John Radcliffe Hospital, University of Oxford, Headley Way, Oxford, OX3 9DU, UK

[2]National Institute of Health Research Oxford Biomedical Research Centre, John Radcliffe Hospital, Headley Way, Oxford, OX3 9DU, UK

### Tables of Free Energies

Three data tables – one for Set1, one for Set2 and one for Set 3 (Table 1) – are included along with the paper. These were produced by analysing the alchemical molecular dynamics simulations and were used in all the analyses in the main body of the paper, as well as the figures in this Supplement. Each table has free energies calculated at 500 different values of the simulation duration, $t$, and hence $\Delta t$ is 0.5ps, 5 ps and 50 ps for Set1, Set2 and Set3, respectively. Each calculation is uniquely identified by the combination of `CALCULATION` and `RUN`. `CALCULATION` only takes one of three values `ti-02`, `ti-05` or `ti-12`. Those labelled `ti-05` contains 10 different values of `RUN` and these are the calculations which were extended from 0.25 ns to 2.5 ns and therefore make up Set2, whilst the `ti-12` contains five separate calculations for the F99Y mutation generated from simulations 25 ns long. Hence, with the exception of F99Y, `dhfr-Set2.csv` only contains `CALCULATION=='ti-05'` whilst `dhfr-Set1.csv` contains both values. The tables not only contain $\Delta\Delta G_{tmp}$ (labelled `ddG_tmp`), but also values of $\Delta\Delta G_{fol}$ (labelled `ddG_fol`, if available) and the individual alchemical free energies used to calculate each. These are labelled with a numerical subscript which refers to the thermodynamic cycle found in Fig. S1. Note that Fig. S1 only contains the labels for $\Delta\Delta G_{tmp}$: those for $\Delta\Delta G_{fol}$ map as follows: `ddG_5` $\rightarrow$ `ddG_8`, `ddG_6[1,2,3]` $\rightarrow$ `ddG_9[1,2,3]` and `ddG_7` $\rightarrow$ `ddG_10`.

[*]published in the Royal Society journal, *Interface Focus*

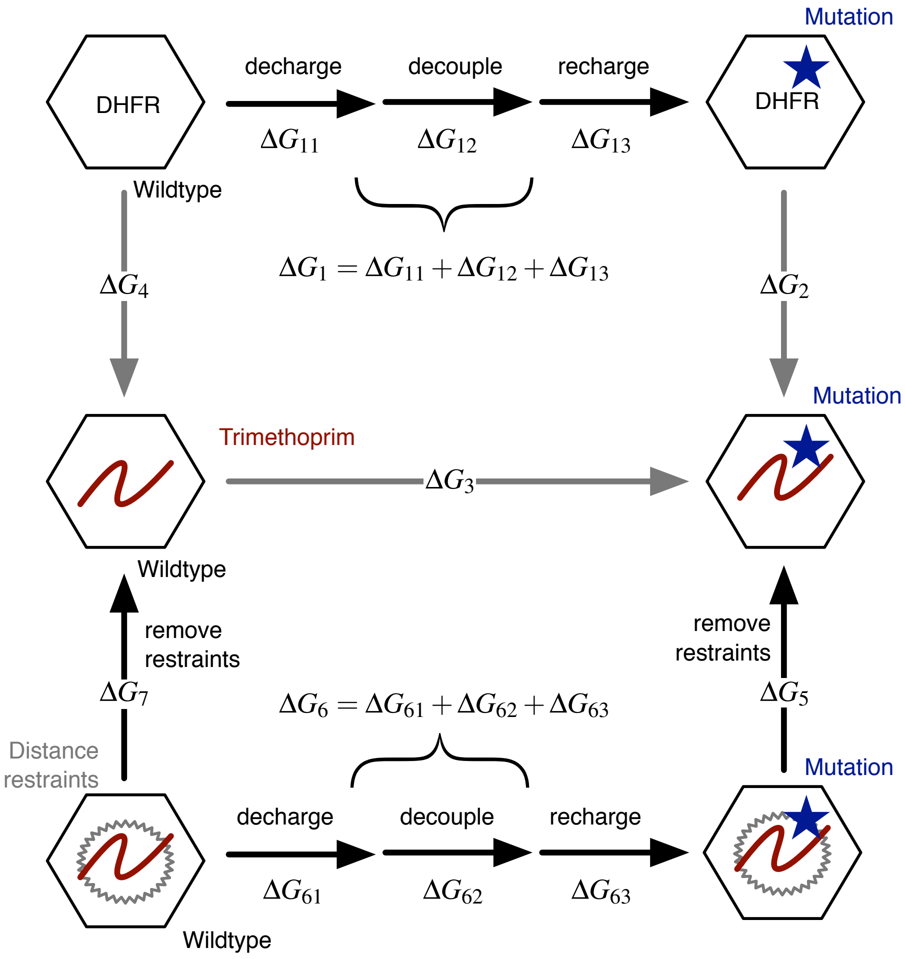


Figure S1: (Related to Equation 1) The thermodynamic cycle used to calculate $\Delta\Delta G_{tmp}$. The nomenclature is used throughout the main body as well as in the included data tables.

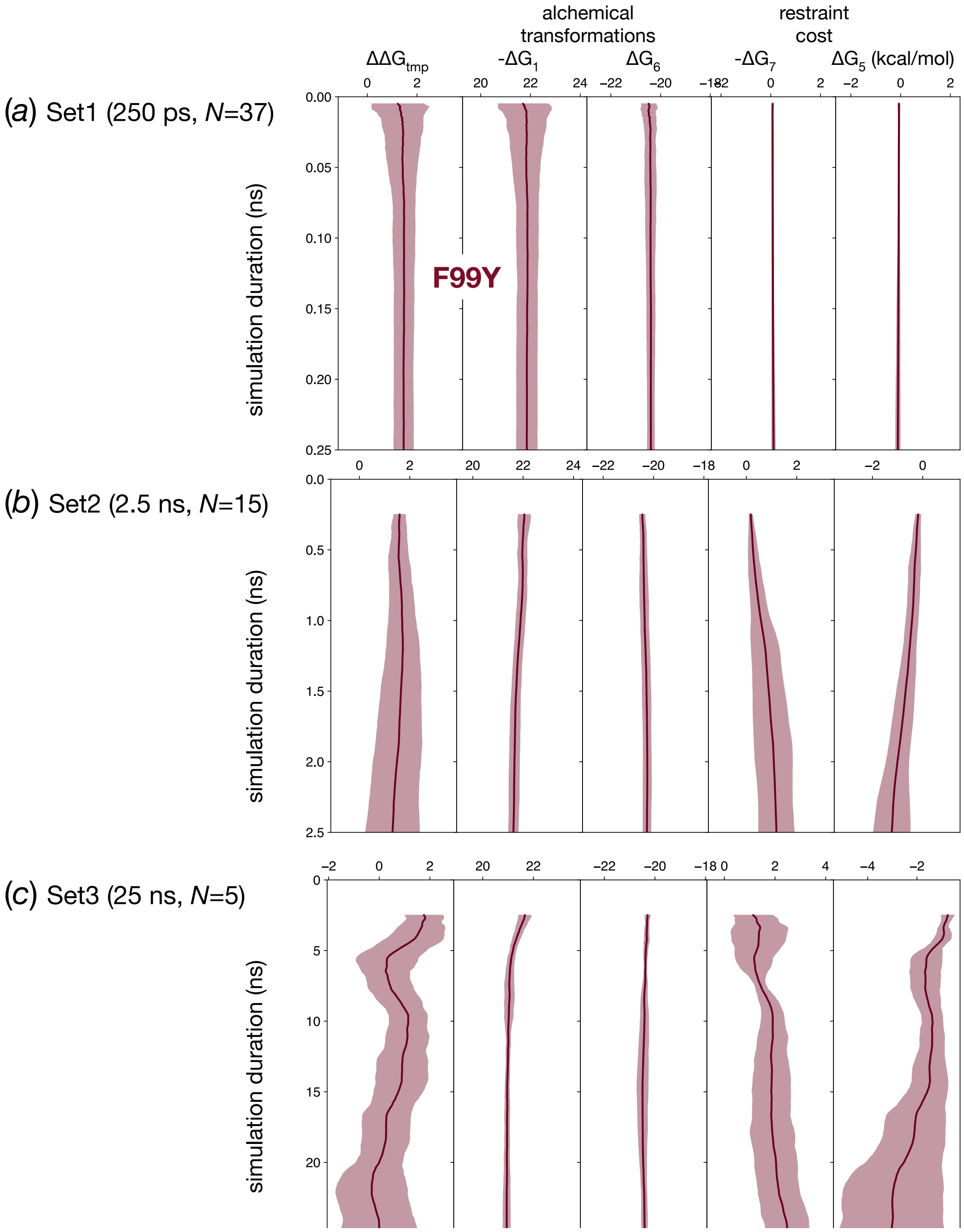


Figure S2: (Related to Figure 2) The mean and variance of the four free energies used to calculate $\Delta\Delta G_{tmp}$ for the F99Y mutation as described by the thermodynamic cycle in Fig. S1 using the varying numbers of simulations belonging to Sets 1, 2 & 3. To aid comparison, the x-axis of each graph covers the same magnitude of energy (5 kcal/mol) and the resulting value of $\Delta\Delta G_{tmp}$ is also shown.

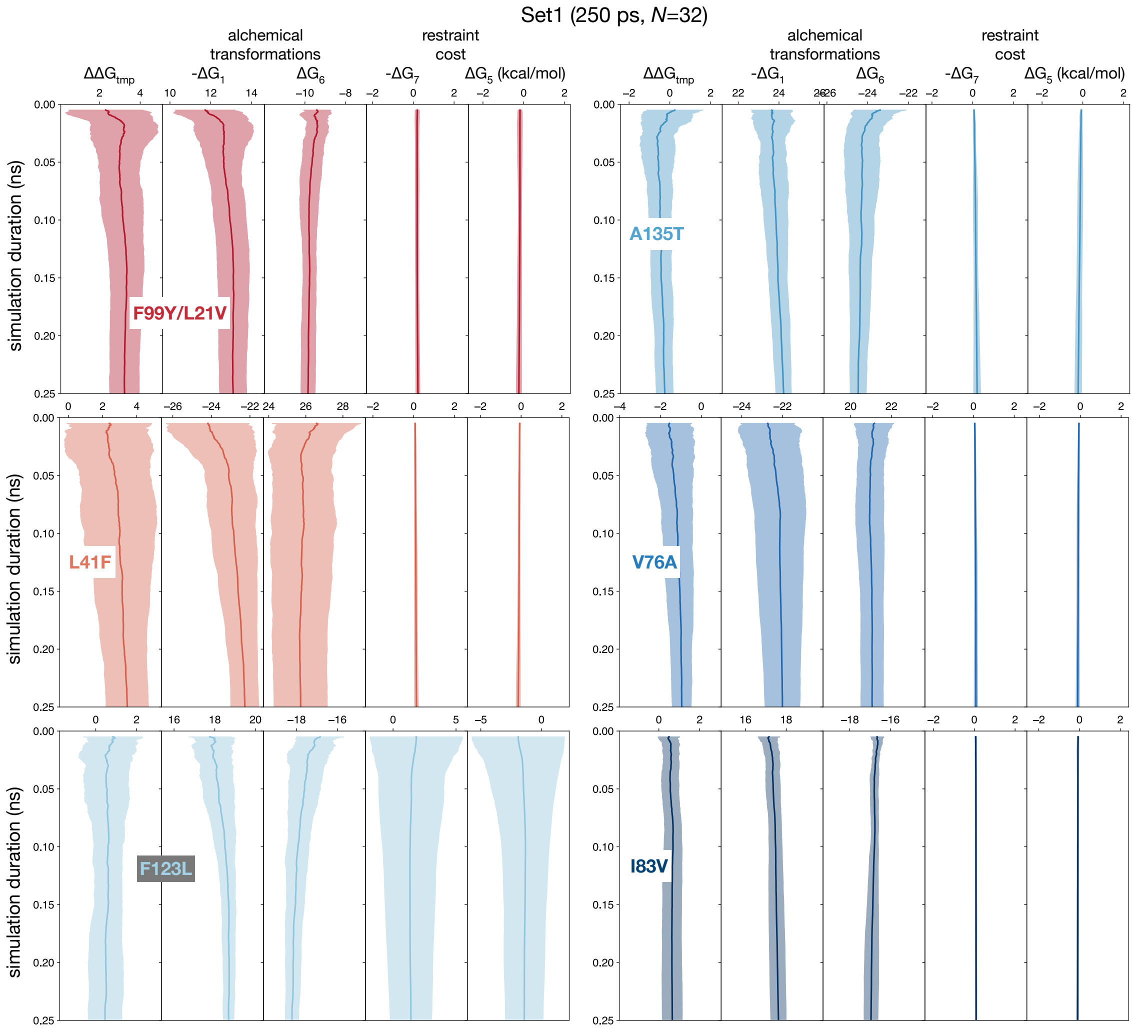


Figure S3: (Related to Figure 3) The mean and variance of the four free energies used to calculate $\Delta\Delta G_{tmp}$ for the other six mutations as described by the thermodynamic cycle in Fig. S1 using the 32 simulations belonging to Set1. To aid comparison, the x-axis of each graph covers the same magnitude of energy (5 kcal/mol) and the resulting value of $\Delta\Delta G_{tmp}$ is also shown.

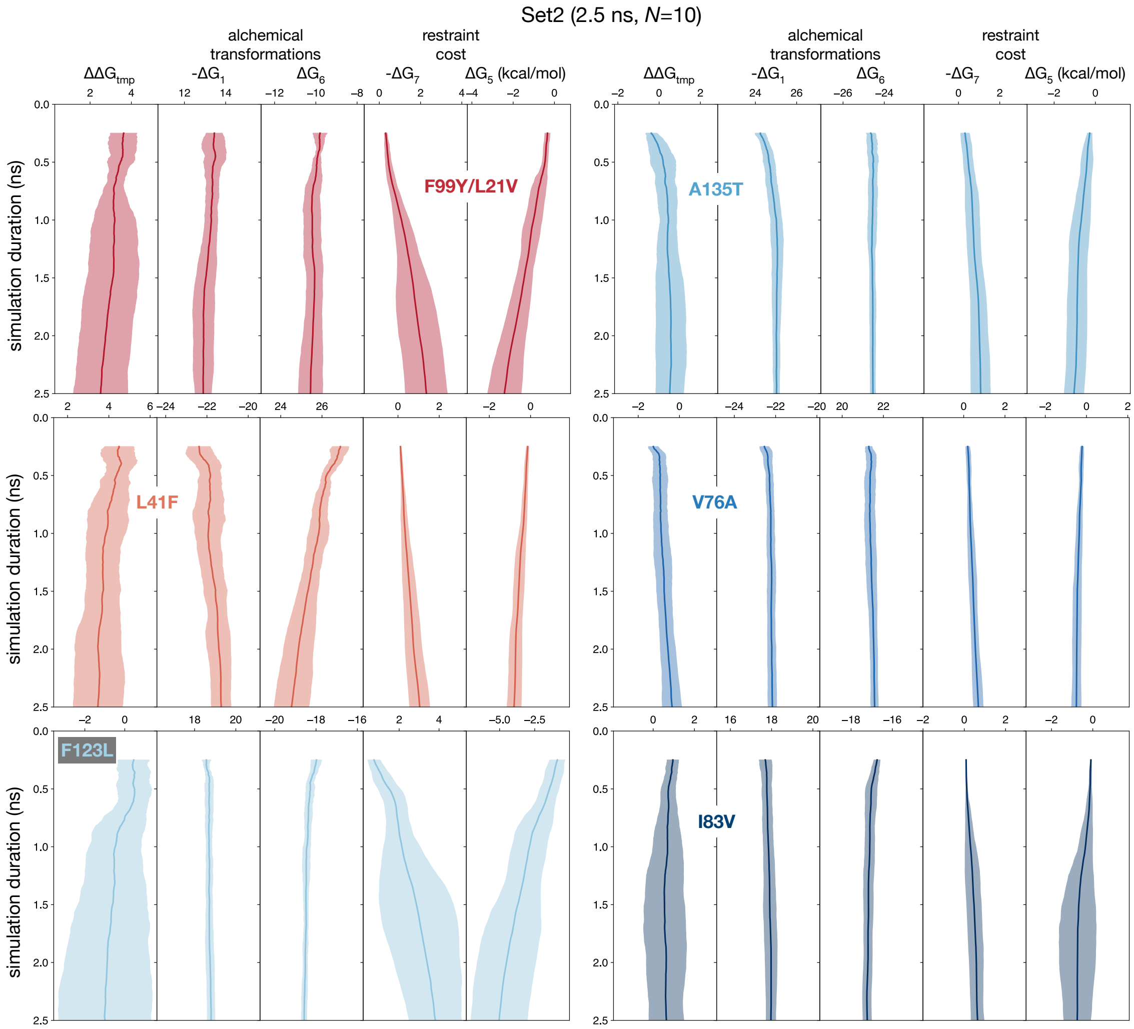


Figure S4: (Related to Figure 3) The mean and variance of the four free energies used to calculate $\Delta\Delta G_{tmp}$ for the other six mutations as described by the thermodynamic cycle in Fig. S1 using the 10 simulations belonging to Set2. To aid comparison, the x-axis of each graph covers the same magnitude of energy (5 kcal/mol) and the resulting value of $\Delta\Delta G_{tmp}$ is also shown.

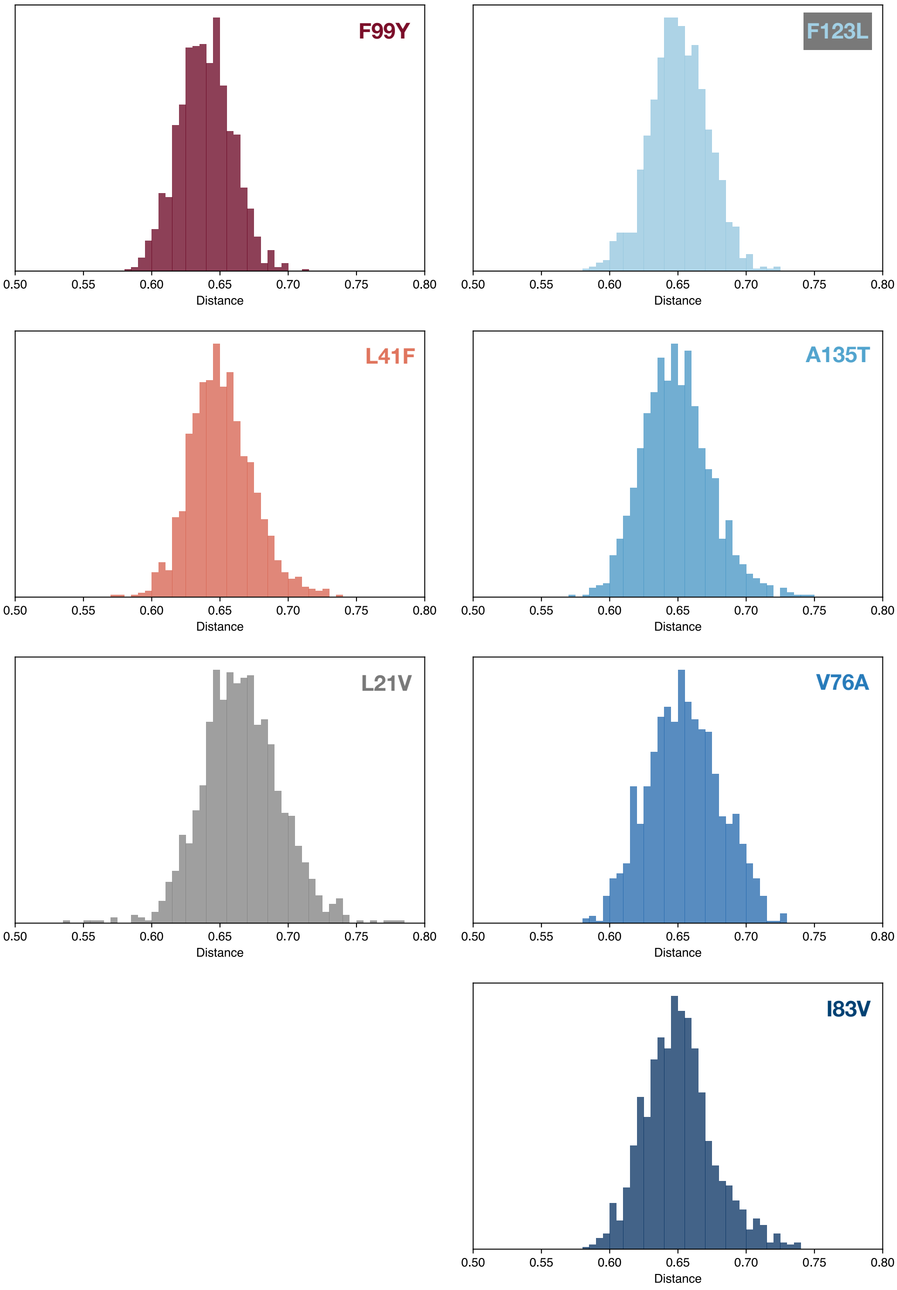


Figure S5: The ligand (trimethoprim) remained bound to the protein in 5× 25 ns unrestrained simulations, suggesting that restraints are not required for these mutations. The distance between the centre of mass of the protein and ligand was calculated for each simulation; these were aggregated for each mutation and a histogram plotted. Note that the Y99L21V simulation was not considered, hence L21V (grey) is shown.

Table S1: Five of the seven amino acid mutations studied were found in a search index of microbial genomes deposited into the European Nucleotide Archive. Only sets of short reads that were assessed as containing at least 80% *S. aureus* were included. Since the index is queried using a *k*-mer, rather than the whole gene, the total number of genomes detected varied across the *dfrB* gene with an average value of 19,200.

| Position | Reference | Mutation | # Genomes | New triplet |
|---|---|---|---|---|
| 12 | caa | Q12R | 9 | cga |
| 14 | gta | V14A | 4 | gca |
| | | V14I | 1 | ata |
| 18 | gaa | E18K | 3 | aaa |
| 21 | tta | L21I | 2 | ata |
| | | L21V | 421 | gta |
| 25 | cta | L25I | 3 | ata |
| 31 | cat | H31N | 15 | aat |
| | | H31Y | 6 | tat |
| 34 | aaa | K34E | 8 | gaa |
| 35 | tta | L35S | 8 | tca |
| | | L35I | 10 | ata |
| 41 | tta | L41F | 1 | ttt |
| | | L41F | 2 | ttc |
| 43 | atg | M43L | 4 | ttg |
| 44 | ggt | G44A | 3 | gct |
| 54 | cca | P54S | 33 | tca |
| | | P54L | 1 | cta |
| | | P54Q | 1 | caa |
| 56 | ccg | P56L | 1 | ctg |
| | | P56S | 12 | tcg |
| | | P56A | 1 | gcg |
| 60 | aat | N60I | 408 | att |
| 62 | gta | V62A | 3 | gca |
| | | V62L | 6 | tta |
| 67 | aca | T67I | 2 | ata |
| | | T67S | 2 | tca |
| | | T67K | 4 | aaa |
| 71 | gta | V71I | 12 | ata |

Continued on next page

Table S1: Five of the seven amino acid mutations studied were found in a search index of microbial genomes deposited into the European Nucleotide Archive. Only sets of short reads that were assessed as containing at least 80% *S. aureus* were included. Since the index is queried using a *k*-mer, rather than the whole gene, the total number of genomes detected varied across the *dfrB* gene with an average value of 19,200.

| Position | Reference | Mutation | # Genomes | New triplet |
|---|---|---|---|---|
| 73 | ggc | G73D | 2357 | gac |
| | | G73S | 1 | agc |
| 76 | gta | V76A | 1412 | gca |
| | | V76L | 1 | tta |
| 78 | cat | H78R | 2 | cgc |
| | | H78P | 1 | ccc |
| | | H78Y | 1 | tat |
| 99 | ttt | F99Y | 137 | tat |
| 103 | att | I103V | 8 | gtt |
| 105 | aaa | K105R | 4 | aga |
| 106 | gtg | V106A | 1 | gcg |
| | | V106M | 7 | atg |
| | | V106E | 1 | gag |
| 125 | cca | P125Q | 4 | caa |
| | | P125L | 2 | cta |
| 131 | gac | D131G | 1 | ggc |
| | | D131N | 1 | aac |
| | | D131Y | 1 | tac |
| 134 | gtt | V134I | 823 | att |
| 135 | gcc | A135T | 6578 | acc |
| | | A135S | 1 | tcc |
| | | A135I | 1 | atc |
| 140 | ggt | G140C | 9 | tgt |
| 144 | gag | E144G | 9 | gga |
| | | E144D | 3 | gat |
| | | E144G | 5 | ggg |
